# Neural Sources Underlying Visual Word Form Processing as Revealed by Steady State Visual Evoked Potentials (SSVEP)

**DOI:** 10.1101/2021.04.16.439729

**Authors:** Fang Wang, Blair Kaneshiro, C. Benjamin Strauber, Lindsey Hasak, Quynh Trang H. Nguyen, Alexandra Yakovleva, Vladimir Y. Vildavski, Anthony M. Norcia, Bruce D. McCandliss

## Abstract

EEG has been central to investigations of the time course of various neural functions underpinning visual word recognition. Recently the steady-state visual evoked potential (SSVEP) paradigm has been increasingly adopted for word recognition studies due to its high signal-to-noise ratio. Such studies, however, have been typically framed around a single source in the left ventral occipitotemporal cortex (vOT). Here, we combine SSVEP recorded from 16 adult native English speakers with a data-driven spatial filtering approach—Reliable Components Analysis (RCA)—to elucidate distinct functional sources with overlapping yet separable time courses and topographies that emerge when contrasting words with pseudofont visual controls. The first component topography was maximal over left vOT regions with an early latency (approximately 180 msec). A second component was maximal over more dorsal parietal regions with a longer latency (approximately 260 msec). Both components consistently emerged across a range of parameter manipulations including changes in the spatial overlap between successive stimuli, and changes in both base and deviation frequency. We then contrasted word-in-nonword and word-in-pseudoword to test the hierarchical processing mechanisms underlying visual word recognition. Results suggest that these hierarchical contrasts fail to evoke a unitary component that might be reasonably associated with lexical access.

## 1 Introduction

Reading is a remarkable aspect of human cognitive development and is essential in everyday life. Through frequent exposure to printed words, visual specialization for letter strings is developed (Maurer et al., 2006), and skilled readers can read around 250 words per minute (Rayner, 1998). Such high reading speed requires fast visual word recognition that is dependent on specialized visual processes and brain sources.

Functional magnetic resonance imaging (fMRI) studies have reliably localized an area of the left lateral ventral occipitotemporal cortex (vOT) that is particularly selective to printed words relative to other visual stimuli such as line drawings (Centanni et al., 2017) and faces (Baker et al., 2007; Dehaene et al., 2005; Dehaene & Cohen, 2011). The sensitivity of the left vOT site to visual words is reproducible across different languages and fonts (Krafnick et al., 2016) and individuals (Dehaene et al., 2010; McCandliss et al., 2003), and is invariant to font, size, case, and even retinal location (Dehaene et al., 2001, 2004).

Numerous fMRI studies have proposed that the vOT follows a hierarchical posterior-to-anterior progression, with posterior regions being more selective to visual word form processing while anterior parts are more weighted to high-level word features (Dehaene et al., 2005; Vinckier et al., 2007). The posterior-to-anterior gradient is accomplished by increasing neuron receptive fields from posterior occipital to anterior temporal regions, and as a result, the sensitivity of neurons hierarchically increases from letter fragments to individual letters, bigrams, trigrams, morphemes, and finally entire word forms (Dehaene et al., 2005). A recent study by Lerma-Usabiaga et al. (2018) suggested that, in addition to sub-regions within the vOT, other regions of the language network (e.g., angular gyrus) are also involved in the rapid identification of word forms by transferring and integrating information from and towards the vOT. Lerma-Usabiaga et al. (2018) further suggested that the posterior area of vOT is structurally connected to the intraparietal sulcus mostly through a bottom-up pathway while the middle/anterior area is connected to other language areas most likely through both feed-forward and -backward connections (see also Price & Devlin (2011)).

In contrast to fMRI with its high spatial resolution, electroencephalography (EEG) can detect text-related brain electrical activity with high temporal resolution. Event-related potentials (ERP) studies have characterized a component that peaks between 150 to 200 ms with an occipito-temporally negative and fronto-centrally positive topography, termed N1 (or N170). This N1 component is typically larger for word and word-like stimuli than for visually controlled symbols (Brem et al., 2006; Maurer et al., 2005, 2006). During development, N1 sensitivity to printed words emerges when children learn print-speech sound correspondences, especially in alphabetic languages (Brem et al., 2010) within the first two years of school reading education (Maurer et al., 2005).

More recently, Steady-State Visual Evoked Potential (SSVEP) paradigm have also been used to investigate visual word recognition due to its high signal-to-noise (SNR) ratio. In contrast to typical ERP approaches demanding long inter-stimulus intervals, the SSVEP paradigm presents a sequence of stimuli at a fast periodic rate (e.g., 10 Hz, 100 ms per item). The presentation of temporally periodic stimuli elicits periodic responses at the predefined stimulation frequency and its harmonics (i.e., integer multiples of the stimulus frequency). Those periodic responses are referred to as SSVEP because they are stable in amplitude and phase over time (Regan, 1966, 1989). Importantly, the SSVEP paradigm can provide high SNR ratio in only a few minutes of stimulation due to its small noise bandwidth. However, the SSVEP paradigm has long been limited to the field of low-level visual perception and attention (for a review, see Norcia et al. 2015). Only recently, this paradigm has been extended to more complex visual stimuli processing, such as objects (Stothart et al., 2017), faces (Alonso-Prieto et al., 2013; Farzin et al., 2012; Liu-Shuang et al., 2014), numerical quantities (Guillaume et al., 2018; Van Rinsveld et al., 2020), text (Yeatman & Norcia, 2016), letters (Barzegaran & Norcia, 2020), and words (Lochy et al., 2015, 2016, 2018, 2020). These SSVEP studies of higher-level processes have used different presentation paradigms including adaptation and “base/deviant” approaches. The adaptation approach has been used to examine whether stimulus presentation locations affect perception, mainly in relation to holistic processing of faces, by comparing upright or inverted faces with either the same or different identity (Rossion et al., 2012). In the “base/deviant” stimulation mode, a sequence of “base” stimuli are presented at a periodic rate (e.g., 6 Hz) with every other image being either an intact image that differs from the base in a particular aspect or a scrambled one (Farzin et al., 2012; Yeatman & Norcia, 2016). For example, a base of 6 Hz alternates with a 3 Hz “deviant” (6/2=3 Hz), which is also called “image alternation” mode. Alternatively, the base stimuli are regularly interspersed with deviant stimuli at a sub-multiple of the base rate that is greater than two (Liu-Shuang et al., 2014; Lochy et al., 2015, 2016, 2018)—for example, a base of 10 Hz with every 5th image (instead of every other image) being a deviant, i.e., deviant frequency is 2 Hz (base frequency 10 Hz divided by 5). However, to our knowledge, no study has directly compared the “image alternation” mode and the mode wherein deviant stimuli are presented at a sub-multiple greater than twice the base rate. Therefore, the current study compared these two modes to determine which of these two modes elicits responses with a higher signal-to-noise ratio. Findings from this research would provide an important consideration relevant to designing future studies of early readers.

Of present interest, two SSVEP studies on text and letter recognition have revealed multiple underlying sources with different temporal dynamics and scalp topography (Barzegaran & Norcia, 2020; Yeatman & Norcia, 2016) either by defining different regions of interest (Yeatman & Norcia, 2016) or employing a spatial filtering approach (Barzegaran & Norcia, 2020). Other SSVEP work has focused on only a single source of word-related processing in the left hemisphere (Lochy et al., 2015, 2016, 2018) by analyzing periodic responses from several pre-selected (literature-based) sensors. In contrast to data analyses of several preselected sensors that reduced the whole map of evoked data to a restricted and typically biased subset (Kilner, 2013), spatial filtering approaches offer a purely data-driven alternative for selecting sensors. These methods compute weighted linear combinations across the full montage of sensors to capture and isolate different neural processes arising from different underlying cortical sources (M. X. Cohen, 2017). A number of linear spatial filters, such as Principal Components Analysis (PCA) and Common Spatial Patterns (CSP) (Blankertz et al., 2008), have been applied to SSVEP data, mainly in the brain-computer interface (BCI) field (e.g., Mohanchandra et al. 2014). In cognitive neuroscience, a spatial filtering technique referred to as Reliable Components Analysis (RCA) has been increasingly used (Barzegaran & Norcia, 2020; Dmochowski et al., 2012, 2015). RCA derives a set of spatial components (i.e., spatial filters operationalized as topographic weights) that maximize across-trial or across-subject correlations (“reliability”) while minimizing noise (“variance”). Specifically, RCA first discovers the optimal spatial filter weighting of the signal, then projects the data through this spatial filter to enable investigation of phase-locked topographic activities and to capture each temporal/topographical source of reliable signal across events and subjects (detailed information described in Dmochowski et al. 2012, 2015).

Moreover, in conjunction with a recently developed RCA approach, Norcia et al. (2020) for the first time estimated the latency of the SSVEP Norcia et al. (2020). This was done by fitting a line through the phases at harmonics with significant responses; the slope of the line is interpreted as the response latency. Latency estimation of component(s) can provide insight into temporal dynamics of different processes located at different sources, extending on the majority of previous SSVEP studies, which have only focused on topographies and amplitudes (Lochy et al., 2015, 2016, 2018).

Employing the spatial filtering component analysis (RCA) approach used in Barzegaran & Norcia (2020) and Norcia et al. (2020), here we reproduced and extended a previous SSVEP study of French word processing (Lochy et al., 2015), with 4-letter English word versus pseudofont comparisons. Our goal was to determine whether multiple sources could be revealed in conditions where only a single source of activity was described. Should this prove to be the case, we were further interested in whether these potential sources could be consistently detected at different stimulus presentation rates and retinal locations.

Specifically, the current study addresses these questions by presenting familiar words interspersed periodically among control stimuli (i.e., pseudofonts) in three different styles: (1) pseudofont base at a presentation frequency of 10 Hz and word deviant at 2 Hz; (2) same presentation rates as in (1) but with stimulus presentation locations jittered around the center of the screen; and (3) pseudofont base at a presentation frequency of 6 Hz and word deviant at 3 Hz. To investigate the hierarchy of visual word recognition, we included two additional stimulus contrasts: word deviant in nonword base, and word deviant in pseudoword base.

Based on the evidence from Lerma-Usabiaga et al. (2018) and Barzegaran & Norcia (2020), we hypothesized that word deviants among pseudofont base stimuli would elicit more than one neural discrimination source when subjected to RCA, producing at least two reliable signal sources with distinct temporal and topographical information. We sought to further investigate whether RCA component topographies and time-courses were specific to particular experimental parameters or whether similar components could be elicited across a wider range of presentation rates and changes in stimulus locations.

Finally, we wished to test a central assumption in previous reports (e.g., Lochy et al. (2015)) based on the notion that hierarchical aspects of visual word processing can be clearly isolated based on progressively specific contrasts of word-in-pseudofont, word-in-nonword, and word-in-pseudoword. Here, RCA provides a novel opportunity to first investigate distinct component topographies elicited from word-in-pseudofont contrasts, and then to investigate the hypothesis that word-in-nonword and word-in-pseudoword contrasts successfully isolate a subset of these sources.

## 2 Methods

### 2.1 Ethics Statement

This research was approved by the Institutional Review Board of Stanford University. All participants delivered written informed consent prior to the study after the experimental protocol was explained.

### 2.2 Participants

Data from 16 right-handed, native English speakers (between 18.1 and 54.9 years old, median age 20.7 years, 7 males) were analyzed in this study. All participants had normal or corrected-to-normal vision and had no reading disabilities. Data from 5 additional non-native English speakers were recorded, but not analyzed here. After the study, each participant received cash compensation.

### 2.3 Stimuli

The study involved four types of stimuli—words (W), pseudofonts (PF), nonwords (NW), and pseudowords (PW)—all comprising 4 elements (letters or pseudoletters). The English words were rendered in the Courier New font. Pseudofont letter strings were rendered from the Brussels Artificial Character Set font (BACS-2, Vidal & Chetail (2017)), mapping between pseudofont glyphs and Courier New word glyphs. Nonwords and pseudowords were also built on an item-by-item basis by reordering the letters of the words: nonwords were unpronounceable, statistically implausible letter string combinations, while pseudowords were pronounceable and well-matched for orthographic properties of intact words (Keuleers & Brysbaert, 2010). Bigram frequencies were matched between words (M (± SD) = 13664 (± 11007)), pseudowords (M (± SD) = 15177 (± 8549)) and nonwords (M(± SD) = 12775 (± 6065)) (*F*(2, 87) < 1, *p* = 0.57). Stimulus parameters are summarized in Table 1, and example stimuli are shown in Figure 1. All words were common monosyllabic singular nouns. The initial and final letters in all words, pseudowords, and nonwords were consonants. Words were chosen to be frequent (average 97.7 per million) with limited orthographic neighbors (average 2.3, range from 0 to 4) according to the Children’s Printed Word Database (Masterson et al., 2010). Words were also chosen with attention to feedforward consistency. All words were fully feedforward consistent based on rime according to the database provided by Ziegler et al. (1997). When averaging across consistency values for each word’s onset, nucleus, and coda in the database provided by Chee et al. (2020), words had an average token feedforward consistency of 0.79. All in all, there were 30 exemplars of each type of stimulus, for 120 exemplars total. All images were 600 × 160 pixels in size, spanning 7.5 (horizontal) by 2 (vertical) degrees of visual angle.

**Figure 1:**
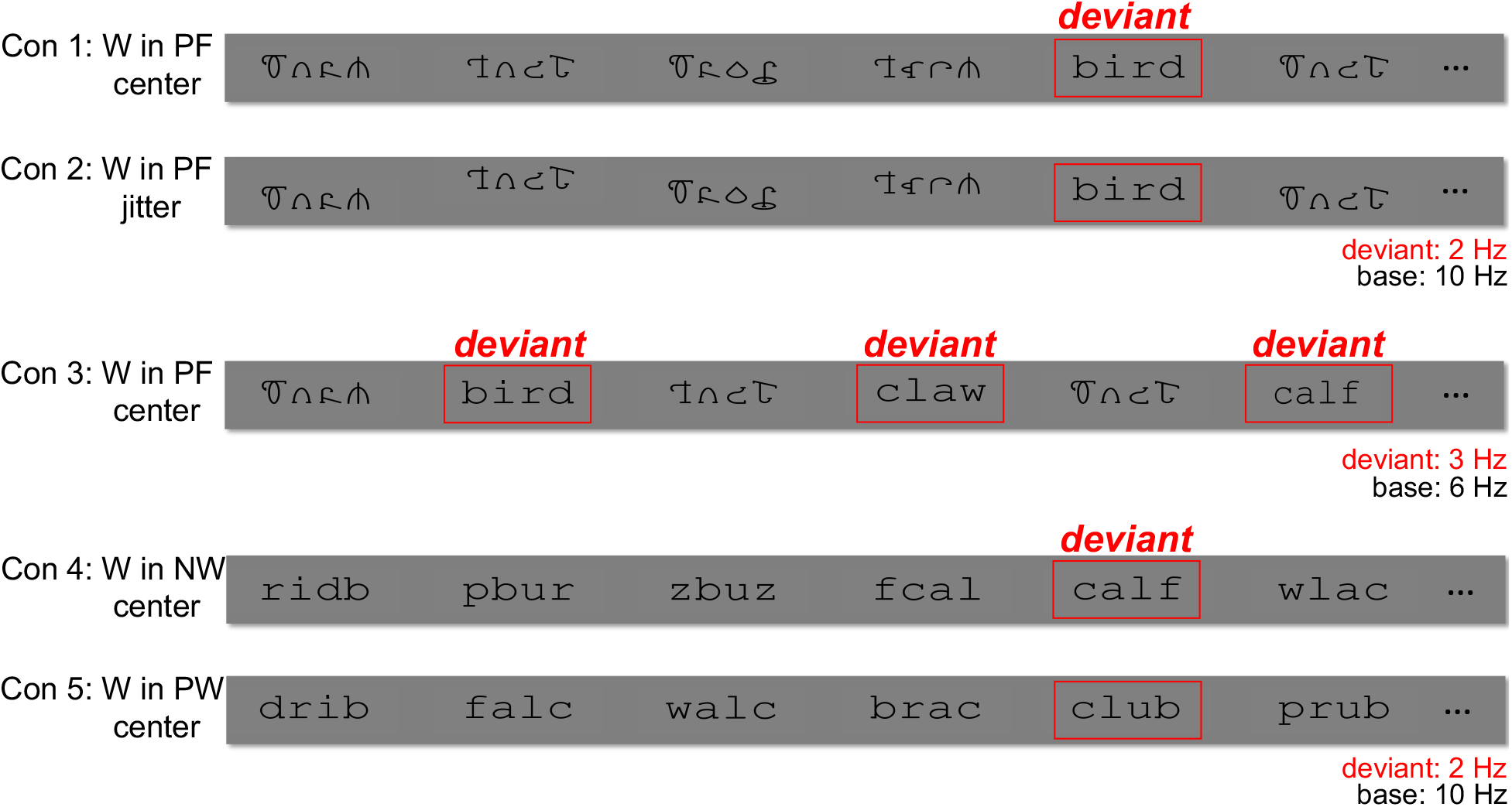
Examples of stimuli presented in the experiment. 2-Hz word deviants were embedded in a 10-Hz stream of pseudofont (W in PF) in conditions 1 and 2. Condition 2 used the same word and pseudofont stimuli as Condition 1, but spatially jittered their location on the monitor. Condition 3 presented the same stimuli used in Condition 1 but with 3-Hz deviant and 6-Hz base frequencies. 2-Hz word deviants were embedded in a 10-Hz stream of nonword (W in NW) and pseudoword (W in PW) base, respectively, in conditions 4 and 5.

**Table 1:**
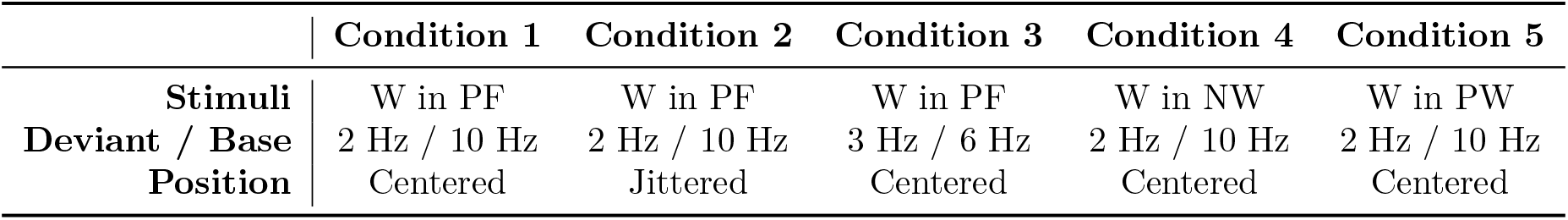
Stimulus conditions. Five stimulus conditions were used to probe different types of processing during word recognition. Conditions 1–3 assessed processing of words (W) relative to pseudofonts (PF), while conditions 4 and 5 assessed processing of words relative to nonwords (NW) and pseudowords (PW), respectively. Word-in-pseudofont contrasts were presented with 2-Hz deviant and 10-Hz base, either centered (condition 1) or jittered (condition 2) around the center of the monitor. The centered word-in-pseudofont contrast was also presented with 3-Hz deviant and 6-Hz base (condition 3). Word-in-nonword (condition 4) and word-in-pseudoword (condition 5) contrasts were presented centered on the screen with frequency rates of 2-Hz deviant and 10-Hz base.

|  | Condition 1 | Condition 2 | Condition 3 | Condition 4 | Condition 5 |
| --- | --- | --- | --- | --- | --- |
| **Stimuli** | W in PF | W in PF | W in PF | W in NW | W in PW |
| **Deviant / Base** | 2 Hz / 10 Hz | 2 Hz / 10 Hz | 3 Hz / 6 Hz | 2 Hz / 10 Hz | 2 Hz / 10 Hz |
| **Position** | Centered | Jittered | Centered | Centered | Centered |

We investigated five experimental conditions. Conditions 1, 2, and 3 involved word deviants embedded in a stream of pseudofont base. For conditions 1 and 2, word deviants were presented at a rate of 2 Hz and were embedded in a 10-Hz stream of pseudofonts. In order to explore the influence of presentation location on word processing, condition 1 stimuli were presented in the center of the screen, while in condition 2, stimulus positions were spatially jittered around the center of the monitor (8 pixels range, visual angle of 0.1 degrees). Moreover, to directly compare the effect of different base/deviant ratios, in condition 3 we used a deviant frequency of 3 Hz (based on the study of Yeatman & Norcia (2016)) and a base frequency of 6 Hz; this involved the same word-in-pseudofont contrast as condition 1, and was presented at the center of the monitor. Finally, conditions 4 and 5 presented word-in-nonword and word-in-pseudoword contrasts, respectively (see also Lochy et al. (2015)). Like condition 1, word deviants were presented at the center of the monitor at 2 Hz, in a base stream of 10 Hz.

### 2.4 Experimental Procedure

Participants were seated in a darkened room 1 m away from the computer monitor. Prior to the experiment, a brief practice session was held to familiarize the participant with the experimental procedure.

Each stimulation sequence started with a blank screen, the duration of which was jittered between 2500 ms and 3500 ms. Then, W deviant stimuli embedded in the stream of base stimuli (PF, NW, or PW) were presented at a rate of 2 Hz (i.e., every 500 ms), with a base rate of 10 Hz (i.e, every 100 ms) in conditions 1, 2, 4, and 5; in condition 3, W deviant were presented at a frequency of 3 Hz (i.e, every 333 ms) with a base rate of 6 Hz (i.e, every 167 ms), during which deviant and base alternated with each other. Thus, a W deviant stimulus was presented every 5 item presentations in conditions 1, 2, 4, and 5, and every 2 items in condition 3. Each condition comprised four trials (each trial lasted for 12 seconds), and was repeated four times, resulting in 16 trials per condition, and 80 trials total in all conditions. 20 trials (5 conditions × 4 trials of each) comprised a block, and the order of trials in each block and all 4 blocks were randomized.

In order to maintain participants’ attention throughout the experiment, a fixation color change task was used. During the recording, the participant continuously fixated on a central cross, which was superimposed over the stimuli of interest and pressed a button whenever they detected that the color of the fixation cross changed from blue to red (2 changes randomly timed per sequence/trial). The color change task was on a “staircase” mode, during which the time of color change flashes became faster as the accuracy increased, or became slower when the accuracy decreased. The whole experiment took around 30 minutes per participant, including breaks between blocks.

### 2.5 EEG Recording and Preprocessing

The 128-sensor EEG were collected with the Electrical Geodesics, Inc. (EGI) system (Tucker, 1993), using a Net Amps 300 amplifier and geodesic sensor net. Data were acquired against Cz reference, at a sampling rate of 500 Hz. Impedances were kept below 50 kΩ. Stimuli were presented using in-house stimulus presentation software. Each recording was bandpass filtered offline (zero-phase filter, 0.3–50 Hz) using Net Station Waveform Tools. The data were then imported into in-house signal processing software for preprocessing. EEG data were re-sampled to 420 Hz to ensure an integer number of time samples per video frame at a frame rate of 60 Hz, as well as an integer number of frames per cycle for the present stimulation frequencies. EEG sensors with more than 15% of samples exceeding a 30 *μ*V amplitude threshold were replaced by an averaged value from six neighboring sensors. The continuous EEG was then re-referenced to average reference (Lehmann & Skrandies, 1980) and segmented into 1-second epochs. Epochs with more than 10% of time samples exceeding a 30 *μ*V noise threshold, or with any time sample exceeding an artifact threshold of (60 *μ*V) (e.g., eye blinks, eye movements, or body movements), were excluded from further analyses on a sensor-by-sensor basis. The EEG signals were filtered in the time domain using Recursive Least Squares (RLS) filters (Tang & Norcia, 1995) tuned to each of the analysis frequencies and converted to complex amplitude values by means of the Fourier transform. Given 1-second data epochs, the resulting frequency resolution was 1 Hz. Complex-valued RLS outputs were decomposed into real and imaginary coefficients for input to the spatial filtering computations, as described below.

### 2.6 Analysis of Behavioral Data

Behavioral responses for the fixation cross color change task served to monitor participants’ attention during EEG recording. We conducted one-way ANOVAs separately for reaction time and accuracy to determine whether participants were highly engaged during the whole experiment.

### 2.7 Analysis of EEG Data

#### 2.7.1 Reliable Components Analysis

Reliable Components Analysis (RCA) is a matrix decomposition technique that derives a set of components that maximizes trial-to-trial covariance relative to within-trial covariance (Dmochowski et al., 2012, 2015). Since response phases of SSVEP are constant over repeated stimulations, RCA uses this trial-to-trial reliability to decompose the entire 128-sensor array into a small number of reliable components (RCs), the activations of which reflect phase-locked activities. Moreover, RCA achieves higher output SNR with a low trial count compared to other spatial filtering approaches such as PCA and CSP (Dmochowski et al., 2015).

Given a sensor-by-feature EEG data matrix (where features could represent e.g., time samples or spectral coefficients), RCA computes linear weightings of sensors—that is, linear spatial filters—through which the resulting projected data exhibit maximal Pearson Product Moment Correlation Coefficients (Pearson, 1896) across neural response trials. The projection of EEG data matrices through spatial filter vectors transforms the data from sensor-by-feature matrices to component-by-feature matrices, with each component representing a linear combination of sensors. For the present study, EEG features are the real and imaginary Fourier coefficients at selected frequencies. As RCA is an eigenvalue decomposition (Dmochowski et al., 2012), it returns multiple components, which are sorted according to “reliability” explained (i.e., the first component, RC1, explains the most reliability in the data). Forward-model projections of the eigenvectors (spatial filter vectors) can be visualized as scalp topographies (Parra et al., 2005). As eigenvectors are known to receive arbitrary signs (Bro et al., 2008), we manually adjusted the signs of the spatial filters of interest based on the maximal correlation between raw sensor data and RCA data. Quantitative comparisons of topographies (e.g., across conditions) were made by correlating these projected weight vectors. Finally, we computed the percentage of reliability explained by individual components using the corresponding eigenvalues, as described by Dmochowski et al. (2015).

#### 2.7.2 RCA Calculations

In order to test whether low-level features were well matched across conditions, we first computed RCA at base frequencies only. Specifically, we input as features the real and imaginary frequency coefficients of the first four harmonics of the base (i.e, 10 Hz, 20 Hz, 30 Hz, 40 Hz for conditions 1, 2, 4 and 5; 6 Hz, 12 Hz, 18 Hz, 24 Hz for condition 3). RCA was computed separately for each stimulus condition.

We next computed RCA at deviant frequencies in order to investigate the processing differences between words and control stimuli (herein pseudofonts, nonwords, and pseudowords). For the deviant analyses, this involved real and imaginary coefficients at the first four harmonics (2 Hz, 4 Hz, 6 Hz, and 8 Hz) in conditions 1, 2, 4, and 5. To explore whether visual word processing can further be consistently detected under different presentation rates, we conducted RCA on data for condition 3. For this, we input frequency coefficients of odd harmonics of the deviant, excluding base harmonics (i.e., 3 Hz, 9 Hz, 15 Hz, 21 Hz—excluding 6 Hz, 12 Hz, 18 Hz, 24 Hz).

To assess the possible role of local adaptation to the stimulus presentation, we also measured responses to word-in-pseudofont using spatially jittered stimuli (condition 2). In comparing RCA results of conditions 1 (word-in-pseudofont with centered presentation location) and 2 (word-in-pseudofont with jittered presentation location), we found the RC topographies of conditions 1 and 2 to be highly correlated (RC1: *r* = 0.99; RC2: *r* = 0.95). Therefore, we subsequently computed RCA on these two conditions together to enable direct quantitative comparison of the projected data in a shared component space.

#### 2.7.3 Analysis of Component-Space Data

For each deviant RCA analysis, we report spatial filter topographies and statistical analysis of the projected data for the first two components returned by RCA. For each component, we analyzed component-space responses at each harmonic input to the spatial filtering calculation. We first projected the sensor-space data through the spatial filter vectors for RCs 1 and 2. The data were averaged across epochs on a per-participant basis, and statistical analyses were performed across the distribution of participants. The distribution of real and imaginary coefficients together at each harmonic formed the basis of a Hotelling’s two-sample t^2^ test (Victor & Mast, 1991) to identify statistically significant responses. We corrected for multiple comparisons using False Discovery Rate (FDR; Benjamini & Yekutieli (2001)) across 8 comparisons (4 harmonics × 2 components per condition).

To test whether phase information was consistent with a single phase lag reflected systematically across harmonics, we fit linear functions through the corresponding phases of successive harmonics, as such a linear relationship would implicate a fixed group delay which can be interpreted as an estimated latency in the SSVEPs (Norcia et al., 2020). At harmonics with significant responses for both RCs (condition 1, conditions 1 and 2 comparison, condition 3), we used the Circular Statistics toolbox (Berens et al., 2009) to compare distributions of RC1 and RC2 phases at those significant harmonics. The results were corrected using FDR across 2 comparisons (2 significant harmonics) for condition 1 and conditions 1 and 2; condition 3 involved no multiple comparisons as only one harmonic was significant. For each of these harmonics, we additionally report each mean RC2-RC1 phase difference in msec.

For each deviant RCA analysis, we present topographic maps of the spatial filtering components, and also visualize the projected data in three ways. First, mean responses are visualized as vectors in the 2D complex plane, with amplitude information represented as vector length, phase information in the angle of the vector relative to 0 radians (counterclockwise from the 3 o’clock direction), and standard errors of the mean as error ellipses. Second, we present bar plots of amplitudes (*μV*) across harmonics, with significant responses (according to adjusted *p_FDR_* values of t^2^ tests of the complex data) indicated with asterisks. Finally, we present phase values (radians) plotted as a function of harmonic; when responses are significant for at least two harmonics, this is accompanied by a line of best fit and slope (latency estimate).

For each base RCA analysis, we report spatial filter topographies and statistical analysis of the projected data for the first component returned by RCA. As with the deviant RCA analysis, we also corrected for multiple comparisons using FDR (Benjamini & Yekutieli, 2001) across 4 comparisons (4 harmonics × 1 component per condition). In contrast to deviant RCA analysis, we visualize the projected data only in bar plots of amplitudes (*μV*) across harmonics, with significant responses (according to adjusted *p_FDR_* values) indicated with asterisks. Phase information and latency estimation are not included here because temporal dynamics are less accurate and less interpretable, especially at high-frequency harmonics (e.g., 30 Hz and 40 Hz) (Cottereau et al., 2011; Norcia et al., 2015).

## 3 Results

### 3.1 Behavioral Results

For the color change detection task, the mean and standard deviation (SD) of accuracy and reaction time across five conditions are summarized in Table 2. Separate one-way ANOVAs indicate that there was no significant difference across conditions in either accuracy (*F*(4, 70) = 0.09, *p* = 0.98) or reaction time (*F*(4, 70) = 0.08, *p* = 0.99). Thus, we concluded that participants were sufficiently engaged throughout the experiment^1^.

**Table 2:** Accuracy and reaction time of responses to cross color change. Values are mean (±SD).

|  | Condition 1 | Condition 2 | Condition 3 | Condition 4 | Condition 5 |
| --- | --- | --- | --- | --- | --- |
| **Stimuli** | W in PF | W in PF | W in PF | W in NW | W in PW |
| **Accuracy(%±SD)** | 95.2(±12.7) | 95.6(±11.4) | 94.0(±12.7) | 96.5(±10.4) | 95.8(±12.0) |
| **Reaction Time (ms±SD)** | 356.5(±45.1) | 350.6(±40.7) | 361.4(±38.6) | 352.6(±37.2) | 356.1(±38.0) |

### 3.2 Base Analyses Results

We performed RCA on responses at harmonics of the base frequency in order to investigate neural activity related to low-level visual processing; results are summarized in Figure 2. Panel A displays topographic visualizations of the spatial filters for maximally correlated (RC1) components (reliability explained: 37.5%). Here, all topographies show maximal weightings over medial occipital areas; correlations among components for conditions 1, 2, 4, and 5 are high (*r* ≥ 0.84), and correlation for condition 3 is lower but still high (*r* = 0.78). The plots in Panel B present projected amplitudes (i.e., projecting data through the spatial filter) in bar plots, with statistically significant responses in the first, second, and fourth harmonics (*p_FDR_* < 0.01, corrected for 4 comparisons) in condition 1 (word-in-pseudofont center); the first harmonic (*p_FDR_* < 0.05) in condition 2 (word-in-pseudofont jitter); all four harmonics (*p_FDR_* < 0.05) in conditions 3 (word-in-pseudofont center slower/alternation) and 4 (word-in-nonword center); and the first three harmonics (*p_FDR_* < 0.05) in condition 5 (word-in-pseudoword center). Amplitude comparisons across conditions showed that there is no significant difference between conditions 1, 2, 4, and 5 (*F*(3, 60) = 2.21, *p* = 0.09), while amplitudes in condition 3 are significantly higher than other conditions (*p* < 0. 05).

**Figure 2:**
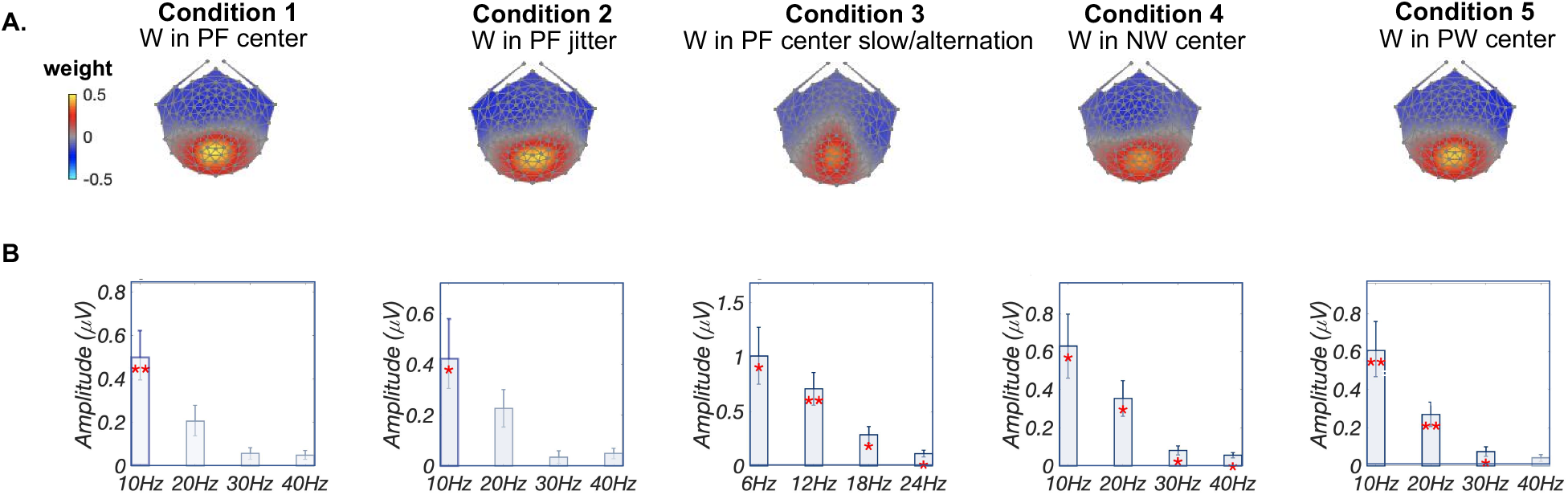
Base Analysis: Low-level visual processing. A: Topographic visualizations of the spatial filters for maximally correlated components (RC1) for all conditions. B: Projected amplitude for each harmonic and condition in bar charts, *: *p_FDR_* < 0.05, **: *p_FDR_* < 0.01.

### 3.3 Deviant Analyses Results

Deviant analyses were conducted to investigate the mechanisms and sources underlying visual word processing. Given increasing evidence supporting segregations within vOT (Lerma-Usabiaga et al., 2018) and the discovery of distinct sources for text and letter recognition (Barzegaran & Norcia, 2020), we hypothesized that multiple sources should be captured during word-related processing (word-in-pseudofont). Due to previous observations of an apparent posterior-to-anterior gradient of responses to visually versus linguistically related sub-regions within vOT (Vinckier et al., 2007; Lerma-Usabiaga et al., 2018), we hypothesized that different condition manipulations (word-in-pseudofont, word-in-nonword, word-in-pseudoword) would evoke different response topographies and phases.

For word deviant responses appearing in a pseudofont base context (word-in-pseudofont, condition 1), the first two reliable components explained the majority (RC1: 34.7%; RC2: 18.7%) of the reliability in the data. As shown in Figure 3A, the topography of the first component (RC1) was maximal over left posterior vOT regions, while the second component (RC2) was distributed over more dorsal parietal regions. Significant signals were present in the first three harmonics of complex-valued data in RC1 and the first two harmonics of RC2 (Figure 3B, see Methods). More detailed amplitude and phase information are presented in Figure 3C. For RC1, the projected data contained significant responses in the first three harmonics (2 Hz, 4 Hz, and 6 Hz; *p_FDR_* < 0.01, corrected for 8 comparisons), while the linear fit across phase distributions for these three harmonics produced a latency estimate of 180.51 ± 0.7 ms. Data projected through the RC2 spatial filter revealed two significant harmonics at 2 Hz and 4 Hz (*p_FDR_* < 0.01) and a longer latency estimate of 261.84 ms (standard error is unavailable when there are two data points). Circular statistics of RC1 and RC2 phase comparisons showed that RC2 phases are significantly longer (2 Hz: 82.9 ms; 4 Hz: 68.1 ms) than RC1 (circular t-test; *p_FDR_* < 0.01 for both 2 Hz and 4 Hz, corrected for 2 comparisons).

**Figure 3:**
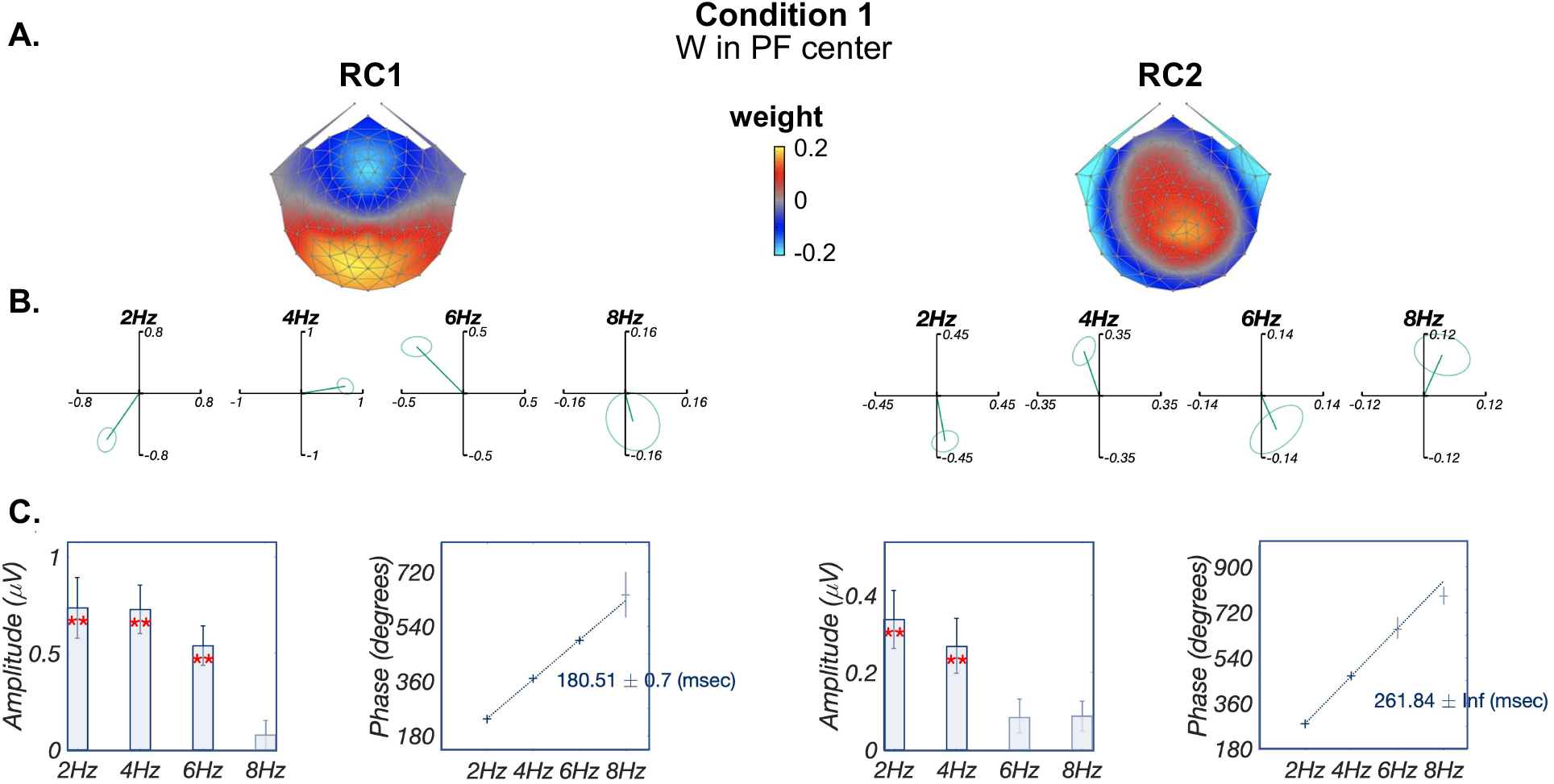
Deviant Analysis: visual word form processing. A: Topographic visualizations of the spatial filters for the first two components (RC1 and RC2); B: Response data in the complex plane, where amplitude information is represented as as the length of the vectors, and phase information in the angle of the vector relative to 0 radians (counterclockwise from 3 o’clock direction), ellipse indicates standard error of the mean for both amplitude and phase; C: Projected amplitude (left) for each harmonic in bar charts, **: *p_FDR_* < 0.01, as well as latency estimations (right) across successful harmonics.

As mentioned in Methods, we found that when word-in-pseudofont stimuli were presented at jittered retinal locations (condition 2), the resulting RC topographies correlated highly with those computed from responses in the centered condition 1 (RC1: *r* = 0.99; RC2: *r* = 0.95). Therefore, in Figure 4 we report RCA results of conditions 1 and 2 in a common component space. As expected, the RC1 and RC2 topographies (Panel A) are similar to those reported when training the spatial filters on condition 1 alone (as presented in Figure 3). The summary plots of the responses in the complex plane (Panel B), show overlapping amplitudes (vector lengths) and phases (vector angles) between these two conditions, especially at significant harmonics (first three in RC1 and first two in RC2). In Panel C response amplitudes did not differ significantly across four harmonics (2 Hz: *p* = 0.70; 4 Hz: *p* = 0.96; 6 Hz: *p* = 0.91; 8 Hz: *p* = 0.65), nor did the derived latency estimations (RC1: 180.42 ms and 173.72 ms for conditions 1 and 2 respectively; RC2: 260.16 ms and 233.32 ms for conditions 1 and 2 respectively). Thus, we did not find evidence that local adaptation is appreciable for these stimuli. Additionally, circular statistics of RC1 and RC2 phase comparisons showed that RC2 phases are significantly longer (2 Hz: 83.2 ms; 4 Hz: 42.2 ms) than RC1 (*p_FDR_* < 0.05 for both 2 Hz and 4 Hz, corrected for 2 comparisons).

**Figure 4:**
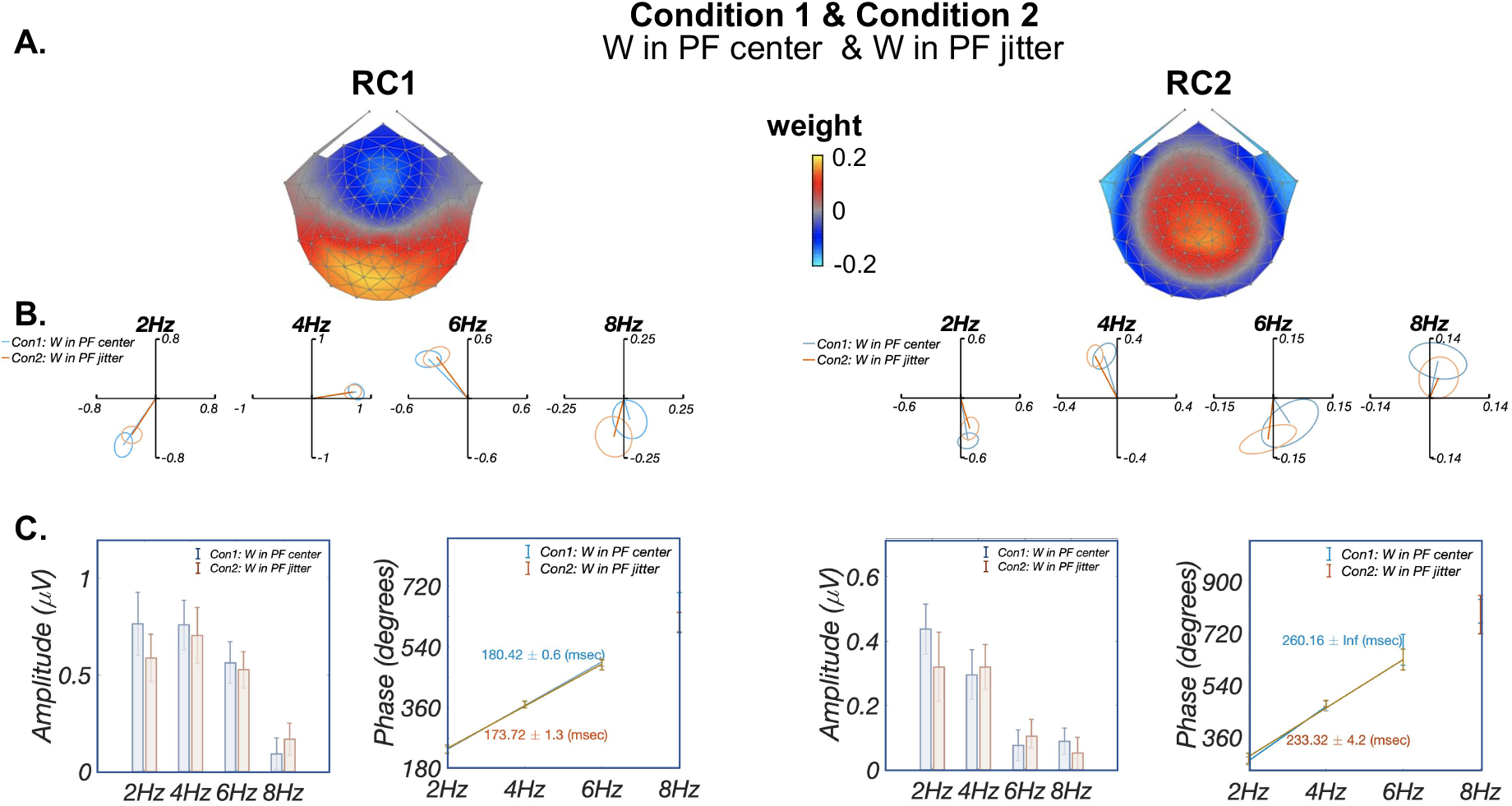
Deviant Analysis: visual word form processing responses are similar irrespective of stimulus location. RCA was trained on response data from conditions 1 and 2 together as their RCs were highly similar when trained separately. A: Topographic visualizations of the spatial filters for the first two components (RC1 and RC2); B: Response data in the complex plane, with condition 1 (W in PF center) in blue and condition 2 (W in PF jitter) in red. Amplitude (vector length) and phase (vector angle, counterclockwise from 0 radians at 3 o’clock direction) overlap across conditions especially at significant harmonics (first three harmonics in RC1 and first two harmonics in RC2). Ellipses indicate standard error of the mean; C: Comparison of projected amplitude (left) and latency estimation (right) between two conditions. Response amplitudes did not differ significantly across the four harmonics (*p* > 0.65), latency estimation derived from phase slopes across harmonics were also similar between conditions (RC1: 180.42 ms and 173.72 ms for conditions 1 and 2 respectively; RC2: 260.16 ms and 233.32 ms for conditions 1 and 2 respectively)

Furthermore, these two RCs were also detected when presenting word-in-pseudofont at a slower alternation presentation rate, with 3 Hz deviant and 6 Hz base (condition 3, Figure 5). Similar to conditions 1 and 2, topographies between conditions 1 and 3 were highly correlated (RC1: *r* = 0.99; RC2: *r* = 0.91). Although it was not possible to estimate the latency, as responses for condition 3 were significant only at the first harmonic (i.e., 3 Hz) for both components (RC1: *p_FDR_* < 0.001; RC2: *p_FDR_* < 0.01, Panel C), circular statistics showed that RC2 phase at 3 Hz is significantly longer (82.0 ms) than RC1 (*p* < 0.001).

**Figure 5:**
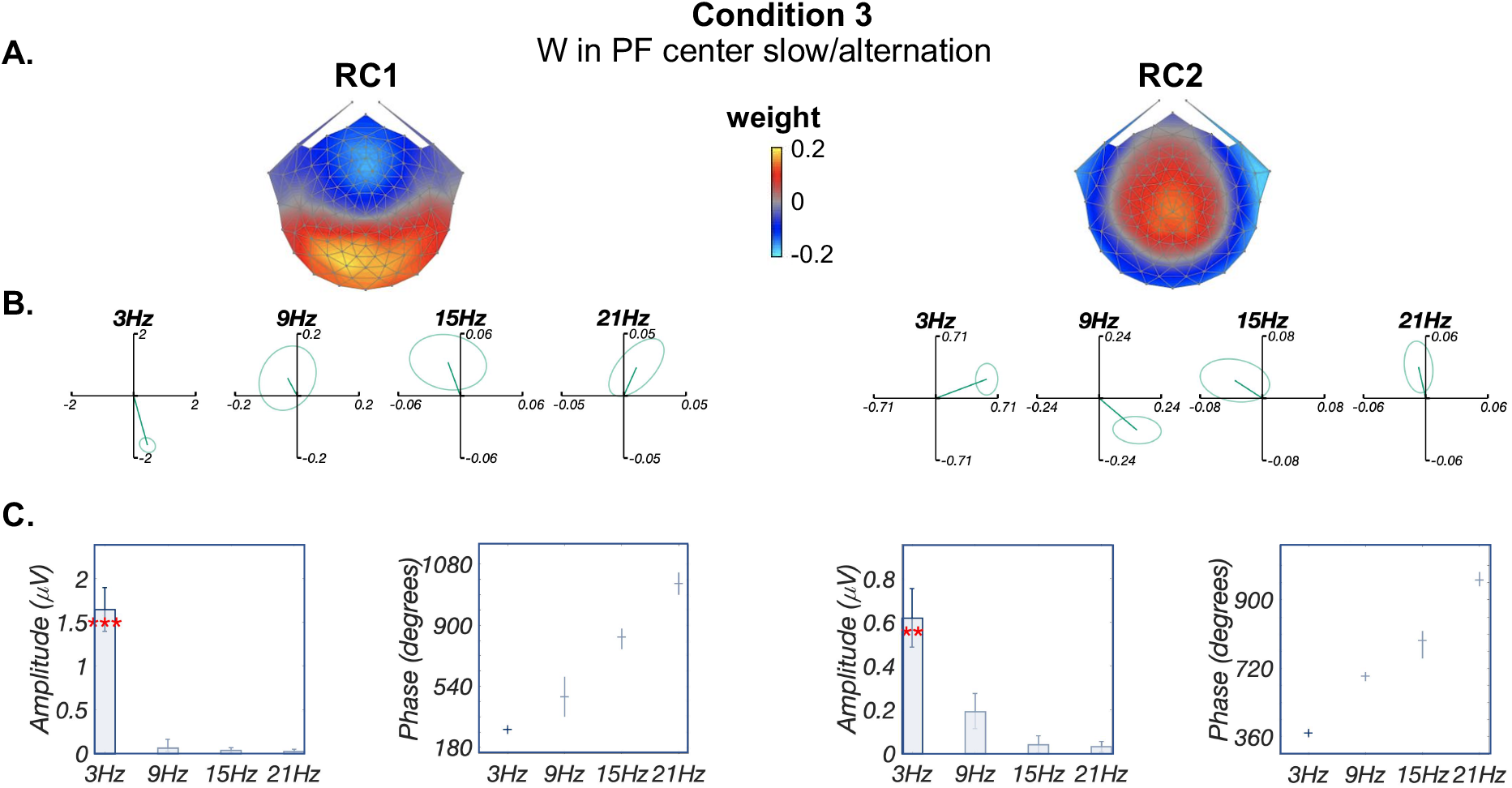
Deviant Analysis: visual word form processing evidenced by changing presentation rates. A: Topographic visualizations of the spatial filters for the first two components (RC1 and RC2); B: Response data in the complex plane, where amplitude information is represented as the length of the vectors, and phase information in the angle of the vector relative to 0 radians (counterclockwise from 3 o’clock direction), ellipse indicates standard error of the mean; C: Projected amplitude for each harmonic in bar charts, **: *p_FDR_* < 0.01, ***:*p_FDR_* < 0.001. Only one significant harmonic prevents us from estimating latency from the phase slope.

Finally, for conditions 4 and 5, word deviants appearing in the other two base contexts (word-in-nonword, word-in-pseudoword) produced components with weaker responses that were associated with distinct topographies, consistent with the hypothesis that each contrast was associated with overlapping yet distinct neural sources (Figure 6). Of note, no more than one significant harmonic is observed for each component: for condition 4 word-in-nonword, only the third harmonic of RC2 is significant (6 Hz, *p_FDR_* < 0.05, corrected for 8 comparisons); for condition 5 word-in-pseudoword no harmonic is significant (*p_FDR_* > 0.14, corrected for 8 comparisons). Due to the lack of significance at multiple harmonics, it was not possible to estimate latencies for these conditions.

**Figure 6:**
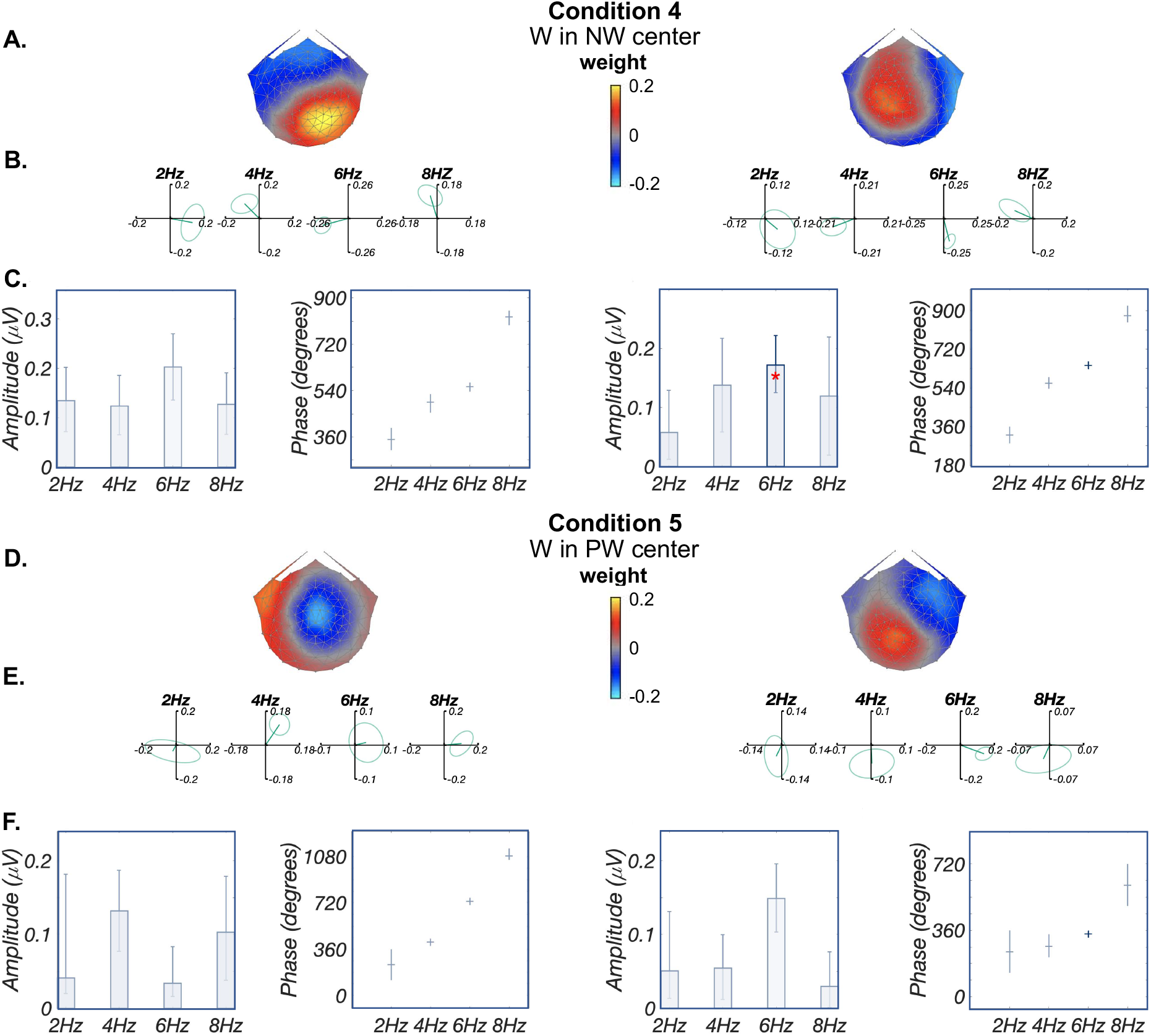
Deviant Analysis: lexical-semantic processing. For condition 4 (W in NW center), A&D: Topographic visualizations of the spatial filters for the first two components (RC1 and RC2) for conditions 4 (W in NW center) and 5 (W in PW center), respectively; B&E: Response data in the complex plane for conditions 4 and 5, respectively. Amplitude information is represented as the length of the vectors, and phase information in the angle of the vector relative to 0 radians (3 o’clock direction), ellipse indicates standard error of the mean. Only the third harmonic has signal for both components in condition 4, while seems the second harmonic for RC1 and the third harmonic for RC2 have signal for condition 5; C&F: Projected amplitude for each harmonic in bar charts, *: *p_FDR_* < 0.05, for condition 4, only the third harmonic is significant in each component; for condition 5, only the third harmonic in RC2 is significant. Having a single significant harmonic prevents us from estimating the phase slope, that’s why no latency estimates were provided.

## 4 Discussion

In this study, we examined the functional and temporal organization of brain sources involved in visual word recognition by employing a data-driven component analysis of Steady-State Visual Evoked Potential (SSVEP) data. We recorded SSVEP word deviant responses appearing in pseudofont base contexts, projecting the multisensor EEG recordings onto single components using Reliable Components Analysis (RCA). Results at the first four harmonics of the base frequency revealed one component centered on medial occipital cortex. Results at harmonics of the deviant frequency revealed two distinct components, with the first component maximal earlier in time over left vOT regions, and the second maximal later in time over dorsal parietal regions. These two components are found to generalize across static versus jittered presentation locations as well as varying rates of stimulation. In addition, distinct topographies were revealed during word deviant responses in the other two base contexts (i.e., pseudowords, nonwords), which—compared with word-in-pseudofont contrast—have different demands of distinguishing words from visual control stimuli.

### 4.1 Low-level visual processing implicates medial occipital areas

RCA analyses of EEG responses at the first four harmonics of the base frequency revealed a maximally reliable component located centered on medial occipital sensors across all conditions. This scalp topography corresponds to expected activations reported in fMRI literature (López-Barroso et al., 2020; Turkeltaub et al., 2003; Szwed et al., 2011) and in EEG source localization studies (Rossion et al., 2003; Proverbio & Adorni, 2009). Medial occipital sensors are directly over early retinotopic visual areas known to support the first stages of visual processing, including of letter strings and objects (Dehaene et al., 2015). Early stages of letter/word and object processing primarily involve low-level visual feature analysis, e.g., luminance, shape, contour, line junctions and letter fragments (Ben-Shachar et al., 2007; Ostwald et al., 2008; Szwed et al., 2011). In our study, basic low-level visual stimulus properties such as spatial frequency, spatial dimensions and the sets of basic line-junction features (Changizi et al., 2006) were well matched across pseudofonts, nonwords and pseudowords base contexts, as well as between base and deviants within each contrast. This may explain comparable responses in terms of amplitudes and topographies across conditions with the same presentation rates. The amplitudes in condition 3 (word-in-pseudofont alternation presentation mode) are higher than other conditions, which may result from higher signal-to-noise ratio in one stimulation cycle of alternation presentation mode Yeatman & Norcia (2016). Nevertheless, consistent responses—in terms of underlying brain sources—across different contrasts, and at different image update rates and retinal locations support that current medial occipital activation reflects low-level visual features processing rather than higher-level word-related processing.

### 4.2 Visual word form processing implicates two distinct sources and processing times

RCA of word deviants in the word-in-pseudofont contrast produced two distinct components with differing latencies. The first component was maximal over ventral occipito-temporal (vOT) regions with slight left lateralization. Phase lag quantification of the first component revealed a linear fit of phases across successive harmonics, providing evidence of a latency estimation around 180 ms.

This 180-ms latency is consistent with the timing of the N170 component revealed in ERP data, which typically peaks around 140-180 ms especially in adults (Eberhard-Moscicka et al., 2015; Maurer et al., 2005). With its characteristic topography over the left occipitotemporal scalp, the N170 is considered to be an electrophysiological correlate of left vOT specialized activation for printed words in fMRI studies, as evidenced by magnetoencephalography (MEG, Hirshorn et al. (2016)), EEG source localization (Brem et al., 2009; Maurer et al., 2005) and simultaneous EEG-fMRI recordings (Pleisch et al., 2019).

Word deviant responses attributed over left-lateralized vOT area in the current study are consistent with numerous fMRI studies showing that the left vOT plays a critical role in fast and efficient visual recognition of words (Cohen et al., 2000, 2002), indicated by selective responses to visually presented words compared with random letter strings and symbols (e.g., pseudofonts) (Dehaene & Cohen, 2011; Rauschecker et al., 2012). This left occipito-temporal activation has also been revealed in SSVEP studies of word recognition (Lochy et al., 2015, 2016). Additionally, using the same RCA approach as in the current study, Barzegaran & Norcia (2020) also reported responses at left vOT cortex when viewing images containing sets of intact versus scrambled letters (Barzegaran & Norcia, 2020). Consistent findings in fMRI, ERP, and SSVEP studies support the reliable involvement of left vOT in processing of words and letters. This specificity of left vOT for words and letters over other visual stimuli is an outcome of literacy acquisition (Dehaene et al., 2015) and emerges rapidly after a short term of reading training (Brem et al., 2010; Chyl et al., 2018; Pleisch et al., 2019), a finding which has also been reported in different writing systems (Bolger et al., 2005; Szwed et al., 2014).

In addition to the first component, we also observed a second RC that was maximal over dorsal parietal regions. A linear fit across phases of successive harmonics demonstrated a latency estimation of around 260 ms, which is consistent with EEG and MEG studies showing phonological effects occurring 250-350 ms after the onset of a visual word (Ashby & Martin, 2008; Grainger et al., 2006; Sliwinska et al., 2012). Finding of a second component is consistent with the increasing evidence showing that visual word recognition is not only limited to the left vOT. Instead, other higher-level linguistic representation areas also support final word recognition (Long et al., 2020; Lerma-Usabiaga et al., 2018; Price & Devlin, 2011). For example, a recent fMRI study showed that, connected with the middle and anterior vOT, other language areas (e.g., the supramarginal, angular gyrus, and inferior frontal gyrus) are responsible for lexical information processing especially for real words (Lerma-Usabiaga et al., 2018). Additionally, anterior vOT is structurally connected to temporo-parietal regions, including the angular gyrus (AG, Booth et al. (2004); Yeatman et al. (2013); Lerma-Usabiaga et al. (2018)), the supramarginal gyri (SMG, Kawabata Duncan et al. (2014); Seghier & Price (2013)) and the superior temporal gyrus (STS, Stevens et al. (2017)). Those areas have been involved in mapping between orthographic, phonological and semantic representations for written words and letter strings (Church et al., 2011; Raij et al., 2000; Van Atteveldt et al., 2004; Vandermosten et al., 2016). In relation to this, an intracranial EEG study suggested the left vOT is involved in at least two distinct stages of word processing: an early stage that is dedicated to gist-level word feature extraction, and a later stage involved in accessing more precise representations of a word by recurrent interactions with higher-level visual and nonvisual regions (Hirshorn et al., 2016). The dorsal parietal brain sources in our study may reflect the later stage of integrating information between anterior vOT (Lerma-Usabiaga et al., 2018) and other language areas (Kay & Yeatman, 2016; Schurz et al., 2014; Woodhead et al., 2014) to enable orthographic-lexical-semantic transformations (Woolnough et al., 2020).

A leading model, the Local Combination Detector (LCD) model (Dehaene et al., 2005), has been proposed to explain the neural mechanisms underlying visual specialization for words. This model suggests that the receptive fields of neurons (detectors) especially in the left vOT progressively increase from posterior to anterior regions. As a result, the sensitivity of neurons increases from familiar letter fragments and fragments combinations to more complex letter strings (e.g, bigrams and morphemes) and whole words. However, given the functional and structural connections between vOT (especially anterior regions) and other language areas mentioned above (i.e., AG, SMG, and STS), a considerable number of studies support the interactive theory in which feed-forward and feed-back processing mechanisms together contribute to final recognition of a word (Dehaene et al., 2005; Dehaene & Cohen, 2011; Dehaene et al., 2015; Long et al., 2020). Specifically, the forward pathway conveys bottom-up progression from early visual cortex (e.g., visual area 4, V4) to vOT, which accumulates inputs about the elementary forms of words (Schurz et al., 2014), and continually from vOT to higher-level linguistic representation areas, which enable integration of orthographic stimuli with phonological and lexical representations (Price & Devlin, 2011). Meanwhile, vOT also receives top-down modulations (backward pathways) from higher-level language regions, which provide (phonology and/or semantic) predictive feedback to the processing of visual attributes.

To summarize, two components were derived in a data-driven fashion from the SSVEP. The first component is located at more left vOT region, while the second component is located more at parietal dorsal area. We speculate that the first component reflects more hierarchical bottom-up forward projections from early visual cortex to posterior vOT, while the second component represents both bottom-up and top-down integration between anterior vOT with other areas of language network. However, these interpretations are speculative without the evidence of source localization data. Thus, more evidence (e.g., source localization and functional connectivity) are needed for verification in future studies.

### 4.3 Reliable Components are invariant to stimulus location and presentation rate

Our results replicated and extended previous work on multiple distinct brain sources involved in different stages of word processing. These findings were further corroborated by presenting stimuli at jittered locations. The response invariance we observe is in agreement with previous studies, in which left vOT activation was identical whether stimuli were presented in the right or left hemifield, suggesting that left vOT activation for words and readable pseudowords depends on language-dependent parameters and not visual features of stimuli (Cohen et al., 2000, 2002). In line with this, Maurer and colleagues (2008) directly compared responses to words and faces under two contexts: blocks that alternated faces and words versus blocks of only faces or words. Results demonstrated that responses to words were consistently left-lateralized and were not manipulated by context in skilled readers. In contrast, context (Maurer et al., 2008) and spatial orientation (Jacques & Rossion, 2007) systematically influenced the degree to which a face is processed. Findings of this kind further suggest that responses to words depend more on linguistic rather than contextual factors (Hauk et al., 2006), which may be driven by the way that words are learned and read (Maurer et al., 2008). Anatomically, it has also been proposed that the visual word form system is homologous to inferotemporal areas in the monkey, where cells are selective to high-level features and invariant to size and position (Cohen et al., 2000). The similarity of the component structure (brain sources) under different presentation rates suggests a fairly broad temporal filter is involved in word vs pseudofont discrimination. Of note, the amplitudes when slowing down the presentation rates are stronger than using higher rates, which sheds light on the choice of the stimulus frequency for studies with children and for examining higher-level visual processes (see also Norcia et al. (2015)).

### 4.4 Hierarchy of visual word processing

Given the word-in-pseudofont contrast results above consistently identified and distinguished two components with distinct topographies and time courses—with the possibility of one being related to processing word visual forms and the other potentially related to higher level integration with language regions—we went on to test whether the hierarchical stimulus contrast approach might also clearly isolate one of these components.

Neither word-in-nonword nor word-in-pseudoword sequences successfully evoke component topographies that resembled either the early or late components of the word-in-pseudofont contrast. Specially, word-in-nonword contrast revealed two components over right occipito-temporal (RC1) and left centro-parietal (RC2) regions, while word-in-pseudoword contrast revealed two components over right centro-frontal (RC1) and left occipito-temporal (RC2) regions. However, component-space neural activations for these two conditions were much weaker and less robust, so caution is needed in their interpretation.

Nevertheless, our weak results in word-in-nonword and word-in-pseudoword contrasts are consistent with a recent study by Barnes et al. (2021). Barnes and colleagues (2021) also used a 4-letter English word vs pseudoword contrast, and the same frequency rates (2 Hz word deviants in 10 Hz pseudoword base) as in our study, results finding that only 4 out of 40 data sets showing a reliable effect between words and pseudowords. While another study by Lochy et al. (2015), which used 5-letter french words and pseudowords with the same 2 Hz/10 Hz rates, revealed response differences between words and pseudowords in 8 out 10 readers. Barnes and colleagues argued that the matching of bigram frequency between words and pseudowords (matched in their study but not in Lochy et al. (2015)) could explain the disparity. Bigram frequency was also matched in our study, which further support Barnes’s argument. Additionally, other factors may also play a role in the disparity, such as number of letters (4 in ours and Barnes et al. (2021), 5 in Lochy et al. (2015)) and number of syllables (monosyllabic in ours and Barnes et al. (2021), monosyllabic and bisyllabic in Lochy et al. (2015)).

Evoked response differences between words and pseudowords and/or nonwords have also been studied previously in ERP studies, but the results have been inconsistent (absent in Araújo et al. (2012); Bentin et al. (1999); Wydell et al. (2003)); present in Eberhard-Moscicka et al. (2015); Kast et al. (2010); McCandliss et al. (1997); Proverbio & Adorni (2009)). Several reasons for the mixed results have been proposed, including but not limited to language transparency, presentation modes and task demands. For example, it was found that the adult N1/left vOT for words is more sensitive to orthographic than lexical and/or semantic contrasts, especially during implicit reading (Bentin et al., 1999; Maurer et al., 2005). Of note, an implicit color detection task was used in the present study, and the pseudowords and nonwords were created by reordering letters that appeared in the words; these factors, in combination, might have led to more difficult discrimination. In addition, words and pseudowords have elicited relatively smaller N170 amplitudes in less transparent English than in German (Maurer 2005). Compared with more transparent languages (e.g., Italian, German and French), English has greater orthographic depth with inconsistent spelling-to-sound correspondence, leading to more ambiguous pronunciations. As a consequence of the inconsistency of mapping letters to sounds, lexical or semantic processing will be less automatic and more demanding (Nosarti et al., 2010), which is even more extreme when reading novel pseudowords and nonwords.

In addition, fast stimulus presentation rates (10 Hz, i.e., 100 ms each item) used in the current study may reduce the involvement of higher-level (e.g., semantic) processes (Vinckier et al., 2007; Lochy et al., 2018) tapped by the word-in-nonword and word-in-pseudoword contrasts, especially during implicit tasks not requiring explicit pronunciation and semantic detection of the stimuli. Lower stimulation rates may be necessary to record SSVEP when discriminating higher-level lexical properties in word-in-nonword and word-in-pseudoword contrasts. By contrast, discrimination of word vs pseudofont is robust over the range of presentation rates we examined. Future studies can examine this issue further by varying stimulation rates over broader ranges or by manipulating task demands.

### 4.5 Future work

In the present study, visual word recognition (W vs PF) was examined using the SSVEP paradigm and a spatial filtering approach, which enabled the identification of robust neural sources supporting distinct levels of word processing within only several minutes’ stimulation. This approach provided a unique extension of existing knowledge on word recognition regarding retinal location and stimulation rate and points to avenues for further investigation of important questions in reading.

First of all, given the short stimulation requirement and robust signal detection due to the high SNR of SSVEP and the spatial filtering approach, the current study points to new possibilities to study individual differences and developmental changes as children learn to read. Children’s reading expertise develops dramatically, especially during the first year(s) of formal reading acquisition (Eberhard-Moscicka et al., 2015; Maurer et al., 2006). The high SNR paradigm used here may allow for more efficient (in terms of less time-consuming and less trials requirement) tracing of the developmental changes of brain circuitry due to increasing reading expertise in children at different stages of learning to read. Additionally, in the course of emerging specialization for printed words, neural component topographies would be expected to be increasingly left-lateralized (Maurer et al., 2005). The RCA approach of detecting multiple, distinct brain sources within the same signal response can increase understanding of how exactly such lateralization is best developed through reading training. Furthermore, high inter-subject variability of vOT print sensitivity was revealed in previous studies (Dehaene-Lambertz et al., 2018; Glezer & Riesenhuber, 2013; Pleisch et al., 2019; Stevens et al., 2017; van de Walle de Ghelcke et al., 2020). Therefore, exploring developmental changes—in terms of activity and topography—at the individual subject level will offer a chance to investigate this phenomenon more precisely and deeply. This approach would be especially relevant to early autistic and dyslexic readers, who face difficulties at the beginning of reading education (Frith & Snowling, 1983) when intervention is thought to be most successful.

Another interesting direction to explore in future research rests upon the accumulating evidence that early visual-orthographic (“perceptual”) and later lexicon-semantic (“lexical”) processing is located at segregated regions within vOT (Lerma-Usabiaga et al., 2018; Stigliani et al., 2015; Vinckier et al., 2007). To our knowledge, different functional components of word recognition have been studied using general contrast of words with pseudofonts and/or consonant strings (Lerma-Usabiaga et al., 2018), but not yet by directly manipulating orthographic regularity and lexical representation(s) separately. Additionally, studies have shown that perceptual tuning of posterior vOT to sublexical and lexical orthographic features develops when reading experience increases (Binder et al., 2006; Zhao et al., 2019). Therefore, isolating perceptual and lexical processing experimentally will be interesting to explore to understand how children’s brains develop specialized visual-orthographic processing.

Last but not least, topographic lateralization and amplitudes of responses to text have been shown to differ when presenting stimuli with different temporal frequencies (Yeatman & Norcia, 2016). In addition, it has been proposed that lower stimulation rates may be necessary to record SSVEPs when studying higher-level visual processes (e.g., faces and words) (Norcia et al., 2015). The relatively high stimulation frequency (i.e., 10 Hz) used in the current study may underestimate effects especially associated with more complex “lexical” contrasts (herein word vs pseudoword and word vs nonword) (Lochy et al., 2018), and this may have an even higher impact when it comes to children. Thus, an important question for future work is to see how children develop specialized visual-orthographic and lexical-semantic processing under lower stimulation rates.

## 5 Conclusion

In conclusion, the present study applied RCA—a data-driven component approach—on SSVEPs, showing that two distinct functional processes with different temporal information underlie visual word form recognition, specifically the word in pseudofont contrast. Those two processes are found to be robust across manipulations of stimulus location as well as stimulation frequency and deviant-base ratio. Moreover, when the same approach was applied to other two contrasts (word in nonword and word in pseudoword), distinct neural sources were found, though their activations were less robust. Our results have provided new evidence that different functional circuits support different stages of word processing. These findings point to novel possibilities toward understanding how visual-orthographic and lexical-semantic processing are best developed through learning experience, which could hold important insights into the acquisition of reading skill.

## Abbreviations

(RCA): Reliable Components Analysis
(RC1): Reliable Component 1
(RC2): Reliable Component 2
(SSVEP): steady-state visual evoked potential

## Footnotes

1 We did find one subject’s accuracy was lower (53.8%) than others. But we still included this participant’s data in our analysis after confirming that this participant showed a cerebral response to the base stimulation (as in Van Rinsveld et al. (2020)). We observed that this participant’s responses to the base rate were not an outlier (i.e., were within one standard deviation) compared with other participants.

## Notes

### Competing Interest Statement

The authors have declared no competing interest.

